# Long-read HiFi Sequencing Correctly Assembles Repetitive *heavy fibroin* Silk Genes in New Moth and Caddisfly Genomes

**DOI:** 10.1101/2022.06.01.494423

**Authors:** Akito Y. Kawahara, Caroline G. Storer, Amanda Markee, Jacqueline Heckenhauer, Ashlyn Powell, David Plotkin, Scott Hotaling, Timothy P. Cleland, Rebecca B. Dikow, Torsten Dikow, Ryoichi B. Kuranishi, Rebeccah Messcher, Steffen U. Pauls, Russell J. Stewart, Koji Tojo, Paul B. Frandsen

## Abstract

Insect silk is an incredibly versatile biomaterial. Lepidoptera and their sister lineage, Trichoptera, display some of the most diverse uses of silk with varying strength, adhesive qualities and elastic properties. It is well known that silk fibroin genes are long (> 20 kb) and have many repetitive motifs. These features make these genes challenging to sequence. Most research thus far has focused on conserved N- and C-terminal regions of fibroin genes because a full comparison of repetitive regions across taxa has not been possible. Using the PacBio Sequel II system and SMRT sequencing, we generated high fidelity (HiFi) long-read genomic and transcriptomic sequences for the Indianmeal moth (*Plodia interpunctella*) and genomic sequences for the caddisfly, *Eubasilissa regina*. Both genomes were highly contiguous (N50 = 9.7 Mbp/32.4 Mbp, L50 = 13/11) and complete (BUSCO Complete = 99.3%/95.2%), with complete and contiguous recovery of silk *heavy fibroin* gene sequences. This study demonstrates that HiFi long-read sequencing can significantly help our understanding of genes with highly contiguous, repetitive regions.

## Background

Many phenotypic traits across the tree of life are controlled by repeat-rich genes [1]. There are many examples, such as antifreeze proteins in fish [2], keratin in mammals, and resilin in insects [1]. Silk is a fundamental biomaterial that is produced by many arthropods, and silk genes are often long (> 20 kb) and contain repetitive motifs [3]. Accurately sequencing through repeat-rich genomic regions is critical to understand how functional genes dictate phenotypes, but research thus far has been unable to quantify these regions. In the case of silk genes, this is essential as these regions control the strength and elasticity properties of silk fibers [4-6].

Lepidoptera (moths and butterflies) and their sister lineage Trichoptera (caddisflies) display some of the most diverse uses of silk from spinning cocoons to prey capture nets and protective armorment [7]. A complete heavy fibroin for the model silkworm moth, *Bombyx mori*, was assembled over two decades ago using BAC libraries [8]. Recently, a combination of nanopore and Illumina sequencing technologies helped generate a full heavy fibroin sequence of *B. mori*, but large regions of the genome remain unassembled [3]. Similarly, we have had similar problems from the Nanopore and Illumina hybrid assemblies in caddisfly genomes [e.g., 9], where we were unable to assemble the complete H-Fibroin genes despite intensive efforts for ∼20 species. In these assemblies, the biggest hindrance was sequencing single strands across large repeat regions and limited illumina polishing due to higher error rates in Nanopore data. The lack of full coverage was largely due to the fact that Nanopore and Illumina sequencing approaches introduce uncertainty for direct inference of function. Therefore, most research thus far has been limited, and focused only on conserved N- and C-terminal regions [e.g., 10]. Complete high-fidelity fully phased fibroin sequences are critical for advancing biomaterials discovery for insect silks.

## Context

We generated HiFi long-read genomic sequences for the Indianmeal moth (*Plodia interpunctella*), and the caddisfly species *Eubasilissa regina*, with the PacBio Sequel II system. Our goal was to recover the area of the genome that has been nearly impossible to sequence due to its repeated regions. We chose these two taxa as they represent two species with very different life histories – *Plodia interpunctella* is an important model organism in Lepidoptera whose larvae feed on a wide variety of grains and stored food products, and secrete large amounts of thin silken webbing at their feeding sites; they also use silk to create a cocoon during pupation [11-12]. *Eubasilissa regina*, on the other hand, is a member of Trichoptera, whose larvae secrete silk in aquatic environments in order to produce protective silk cases made of broader leaf pieces from deciduous trees, cut to size [13]. These new resources not only expand our knowledge of a primary silk gene in Lepidoptera and Trichoptera, but also contribute new, high-quality genomic resources for aquatic insects and arthropods which have thus far been underrepresented in genome biology [14-16].

## Methods

### Sample information and sequencing

A single adult specimen of each species was sampled for inclusion in the present study. For *P. interpunctella*, we used a specimen from the PiW3 colony line at the USDA lab (1600 SW 23 Dr. Gainesville, FL, USA), and its entire body was used for extraction, given its small size. For *E. regina*, a wild-caught female adult specimen (#AK0WP01) from Enzan, Yamanashi, Japan (N35°43’24” E138°50’33”, elevation ∼4,840 ft), originally deposited in the Smithsonian Institution, National Museum of Natural History (USNMENT01414923), was used. The head and thorax were macerated and DNA was extracted. The remainder of the body is preserved as a frozen tissue sample in the lab of PRB at BYU. Both specimens were flash frozen in LN2 and DNA was extracted using Quick-DNA HMW MagBead Kit (Zymo Research). Extractions with at least 1 μg of high-molecular weight (> 40kb) were sheared and the BluePippin system (Sage Science, Beverly, MA, USA) was used to collect fractions containing 15 kb fragments for library preparation. Sequencing libraries were prepared for each species using the SMRTbell Express Template Prep Kit 2.0 (PacBio, Menlo Park, CA, USA) and following the ultra-low protocol. All sequencing was performed using the PacBio Sequel II system. For *P. interpunctella*, the genomic library was sequenced on a single 8M SMRTcell and *E. regina* was sequenced on three 8M SMRTcells, all with 30 hour movie times. For the *P. interpunctella* Iso-seq transcriptome, RNA was extracted using TRIzol (Invitrogen) from freshly dissected silk glands of caterpillars and following manufacturer’s protocol. This species has relatively small body size compared to other Lepidoptera, so we waited until caterpillars reached their maximum size (during the fifth instar) before dissection, in order to maximize yield. Sequencing libraries were prepared following the PacBio IsoSeq Express 2.0 Workflow and using the NEBNext Single Cell/Low Input cDNA Synthesis & Amplification Module for the SMRTbell Express Template Prep Kit 2.0. The resulting library was multiplexed and sequenced on a single Sequel II PacBio SMRT cell for 30 hrs. Library preparation and sequencing was carried out at DNA Sequencing Center at Brigham Young University (Provo, UT, USA).

Genomic HiFi reads were generated by circular consensus sequencing (CCS) where consensus sequences have three or more passes with quality values equal to or greater than 20, from the subreads.bam files and using pbccs tool (v.6.0.0) in the *pbbioconda* package (https://github.com/PacificBiosciences/pbbioconda). Using the same *pbbioconda* package and the Iso-seq v3 tools, high quality (> Q30) transcripts were generated from HiFi read clustering without polishing.

### Genome size estimations and genome profiling

Estimation of genome characteristics such as size, heterozygosity, and repetitiveness were conducted using a *k-*mer distribution–based approach. After counting *k-*mers with K-Mer Counter (KMC) v.3.1.1 and a *k-*mer length of 21 (-m 21), we generated a histogram of *k-*mer frequencies with KMC transform histogram [17]. We then generated genome *k-mer* profiles on the *k-*mer count histogram using the GenomeScope 2.0 web tool [18] with the *k-*mer length set to 21 and the ploidy set to 2.

### Sequence assembly and analysis

For both genomes, reads were then assembled into contigs using the assembler Hifiasm v0.13-r307 with aggressive duplicate purging enabled (option −l 2) [19]. The primary contig assembly was used for all downstream analyses. Genome contiguity was measured using assembly_stats.py [20] and genome completeness was determined using Busco v.5.2.2 [21] and the obd10 reference Endopterygota. Contamination in the genome was assessed by creating Taxon-annotated GC-coverage (TAGC) plots using BlobTools v1.0 [22]. First, assemblies were indexed using *samtools faidx* then HiFi reads were mapped back to the indexed assemblies using minimap2 [23] with *-ax asm20*. The resulting bam files were sorted with *samtools sort*. Taxonomic assignment was performed via Megablast and using the NCBI nucleotide database with parameters -outfmt 6 qseqid staxids bitscore std’ -max_target_seqs 1 -max_hsps 1-e value 1e-25. BlobPlots were created by making a blobtools database from the assembly file, blast results, and mapping results using *blobtools create* and plots were created using *blobtools plot*.

### Genome statistics

All samples, raw sequence reads, and assemblies were deposited to GenBank (Table 1). We generated 35.7 Gbp (41x coverage) and 15.7 Gbp (44x coverage) of PacBio HiFi sequence for *E. regina* and *P. interpunctella*, respectively. We assembled those reads into two contiguous genome assemblies. The assembly for *E. regina* has the highest contig N50 of any Trichoptera genome assembly to date. It contains 123 contigs, a contig N50 of 32.4 Mbp, GC content of 32.68%, and a total length of 917,780,411□bp. GenomeScope 2.0 estimated a genome size of 854,331,742 bp with 75.3% unique sequence (http://genomescope.org/genomescope2.0/analysis.php?code=ghDHLpAQUkIKK4e5yH88). Despite recent analyses showing no evidence of whole-genome duplication in caddisflies (Heckenhauer et al. 2022), the findings in this study may be an indication of tetraploidy. Future research should be done to further examine these patterns.

**Table 1.**
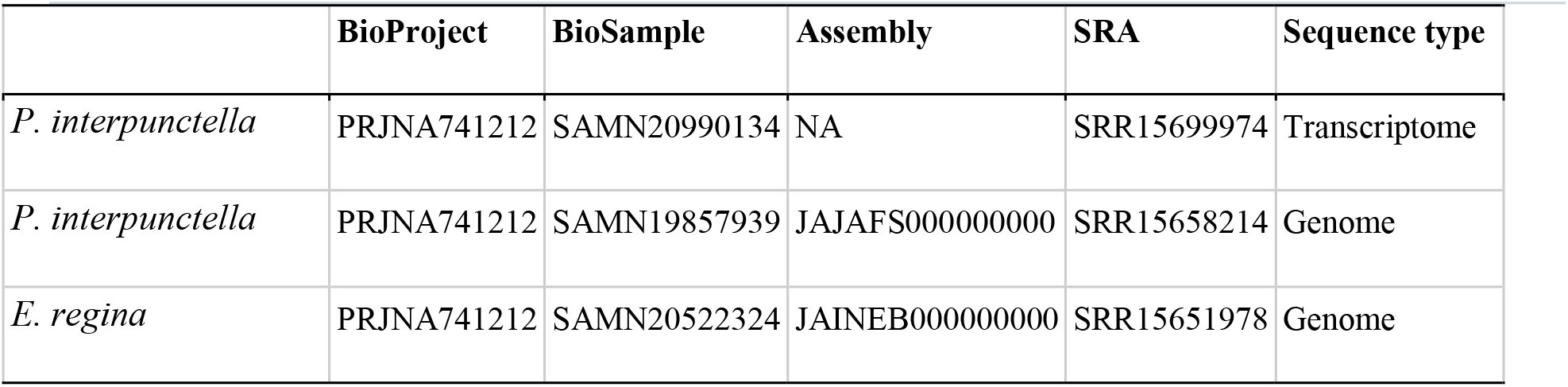
Specimen accession and data type information.

The *P. interpunctella* assembly represents a substantial improvement to existing, publicly available genome assemblies (Table 2). After contaminated contigs were removed (three contigs contaminated with *Wolbachia* were identified), the resulting assembly comprises 118 contigs with a cumulative length of 300,731,903□bp. It exhibits a contig N50 of 9.7Mbp and a GC content of 35.41%. The genome size estimated by GenomeScope 2.0 was 275,458,564 bp with 87.1% unique sequence (http://genomescope.org/genomescope2.0/analysis.php?code=96nVnnk42W5nlBWIfHFj).

**Table 2.** Assembly genome stats for the species sampled in this study.

|  | *P. interpunctella* | *E. regina* | *P. interpunctella* | *P. interpunctella* |
| --- | --- | --- | --- | --- |
| Reference | This study | This study | GCA_001368715.1 | GCA_900182495.1 |
| Platform | PacBio Sequel II | PacBio Sequel II | Illumina MiSeq/HiSeq | Illumina MiSeq/HiSeq |
| Coverage | 44x | 41x | 100x | 50x |
| Total ungapped length | 300,731,903 | 917,780,411 | 364,621,386 | 364,623,808 |
| Total gapped length | NA | NA | 382,235,502 | 381,952,380 |
| Number of scaffolds | NA | NA | 7,743 | 10,542 |
| Scaffold N50 | NA | NA | 5,094,612 | 1,270,674 |
| Scaffold L50 | NA | NA | 23 | 75 |
| Number of contigs | 118 | 123 | 17,717 | 17,725 |
| Contig N50 | 9,707,027 | 32,427,664 | 302,097 | 298,497 |
| Contig L50 | 13 | 11 | 314 | 319 |
| GC content | 35.41% | 32.68% | 35.1% | 35.1% |
| Shortest Contig | 452 | 15,452 | 258 | 258 |
| Longest Contig | 13,555,736 | 57,864,696 | 2,314,344 | 2,314,344 |
| Median Contig | 161,724 | 36,760 | 1,714 | 1,719 |
| Mean Contig | 2,548,575 | 7,401,455 | 20,580 | 20,571 |

**Table 3.** Genome completeness by sample studied. Values shown are BUSCO scores for the Endopterygota ODB10 data set.

|  | *P.*<br>*interpunctella* | *E. regina* | *P. interpunctella* | *P. interpunctella* |
| --- | --- | --- | --- | --- |
|  | This study | This study | GCA_001368715 | GCA_900182495 |
| Complete BUSCOs | 2110 | 2021 | 2103 | 2105 |
| Complete and single-copy | 2097 | 2013 | 2074 | 2077 |
| Complete and duplicated | 13 | 14 | 29 | 28 |
| Fragmented | 5 | 63 | 10 | 8 |
| Missing | 9 | 34 | 11 | 11 |
| Total groups searched | 2124 | 2124 | 2124 | 2124 |
| % complete | 99.3 | 95.2 | 99.0 | 99.1 |

### Heavy fibroin gene annotation

We extracted *heavy fibroin* (*H-fibroin*) silk genes from both the *P. interpunctella* and *E. regina* assemblies. For *P. interpunctella*, we also searched existing, short-read based assemblies. We downloaded two short-read based genome assemblies for *P. interpunctella*, GCA_001368715.1 and GCA_900182495.1 from NCBI (https://www.ncbi.nlm.nih.gov/). Since the internal region of *H-fibroin* is known to be repetitive, the more conserved N- and C-termini amino acids were blasted against the genomes with tblastn. For *P. interpunctella*, we used the terminal sequences published in [24] and for *E. regina*, we used the terminal sequences published in [5]. We then extracted the sequences and 500 bps of flanking regions from the assembly and annotated them using Augustus v.3.3.2 [25]. Spurious introns (those that did not affect reading frames and were not supported by transcript evidence) were manually removed. Annotated sequences are provided in the *Gigascience* GigaDB repository [26].

We recovered full-length *H-fibroin* sequences in both genomes. To our knowledge, the only other previously published full-length lepidopteran *H-fibroin* sequence was from a BAC library-based sequence of the model organism, *B. mori*. We compared our assembly of the *P. interpunctella H-fibroin* sequence with that from a previously published Illumina-based genome assembly of the same species (Table 2). Where the Illumina-based assembly only recovered the conserved terminal regions and a small number of repetitive elements, our assembly recovered the full-length gene, including the full complement of repetitive motifs (Figures 1, 2). Specifically, the *P. interpunctella* genome had a *H-fibroin* sequence that was 14,866 bp (whole gene with introns; 4,714 AA), and a molecular weight of 413,334.41 Da. For *E. regina*, we recovered the full-length sequence of *H-fibroin*, which was 25,250 bp (whole gene with introns; 8,386 AA), and a molecular weight of 815,864.95 Da, with repeated regions (Figure 3). The recovery of this *H-fibroin* sequence marks the third complete, published *H-fibroin* sequence in Trichoptera [27-28]. Our work shows that high quality, long-read sequencing can be used to successfully assemble difficult regions of non-model organisms without the use of expensive and tedious BAC methods. While our study is focused on the repetitive silk gene, *H-fibroin*, these results likely extend to other long, repetitive proteins that have previously proven difficult to assemble.

**Figure 1.**
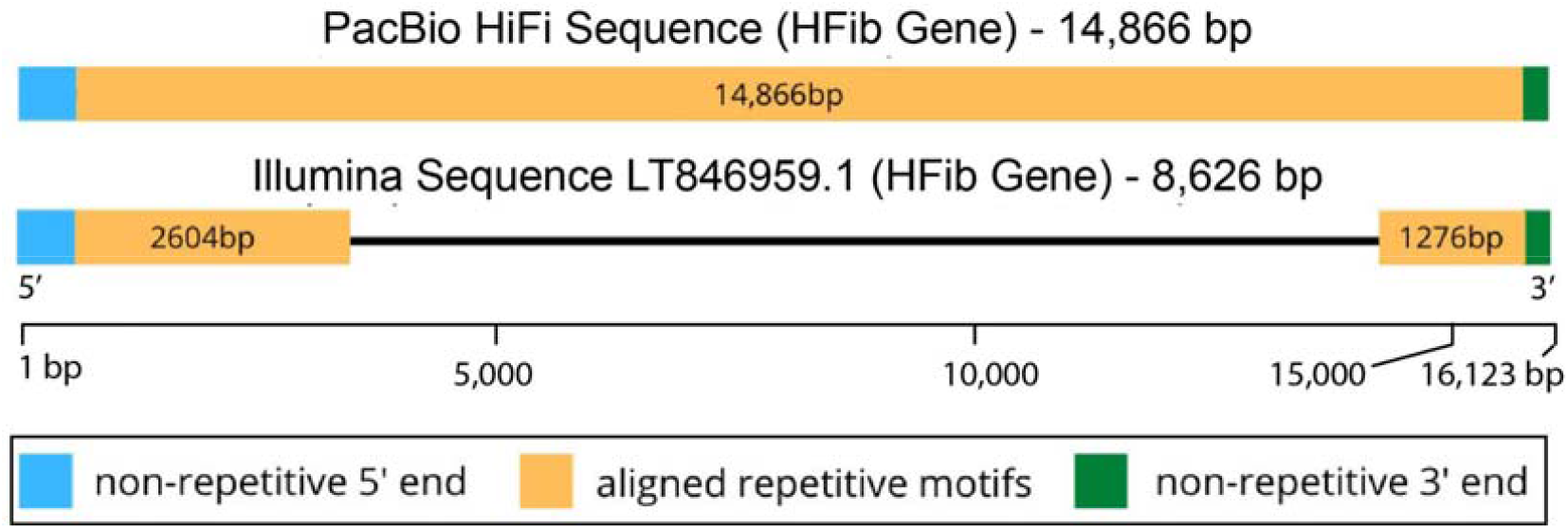
Length of assembled heavy fibroin (HFib) gene with two approaches (HiFi, top; Illumina, bottom) for *P. interpunctella*. In the HiFi genome, we were able to recover the entire length of sequence, but in the latter we were unable to assemble the genome through the repetitive region.

**Figure 2.**
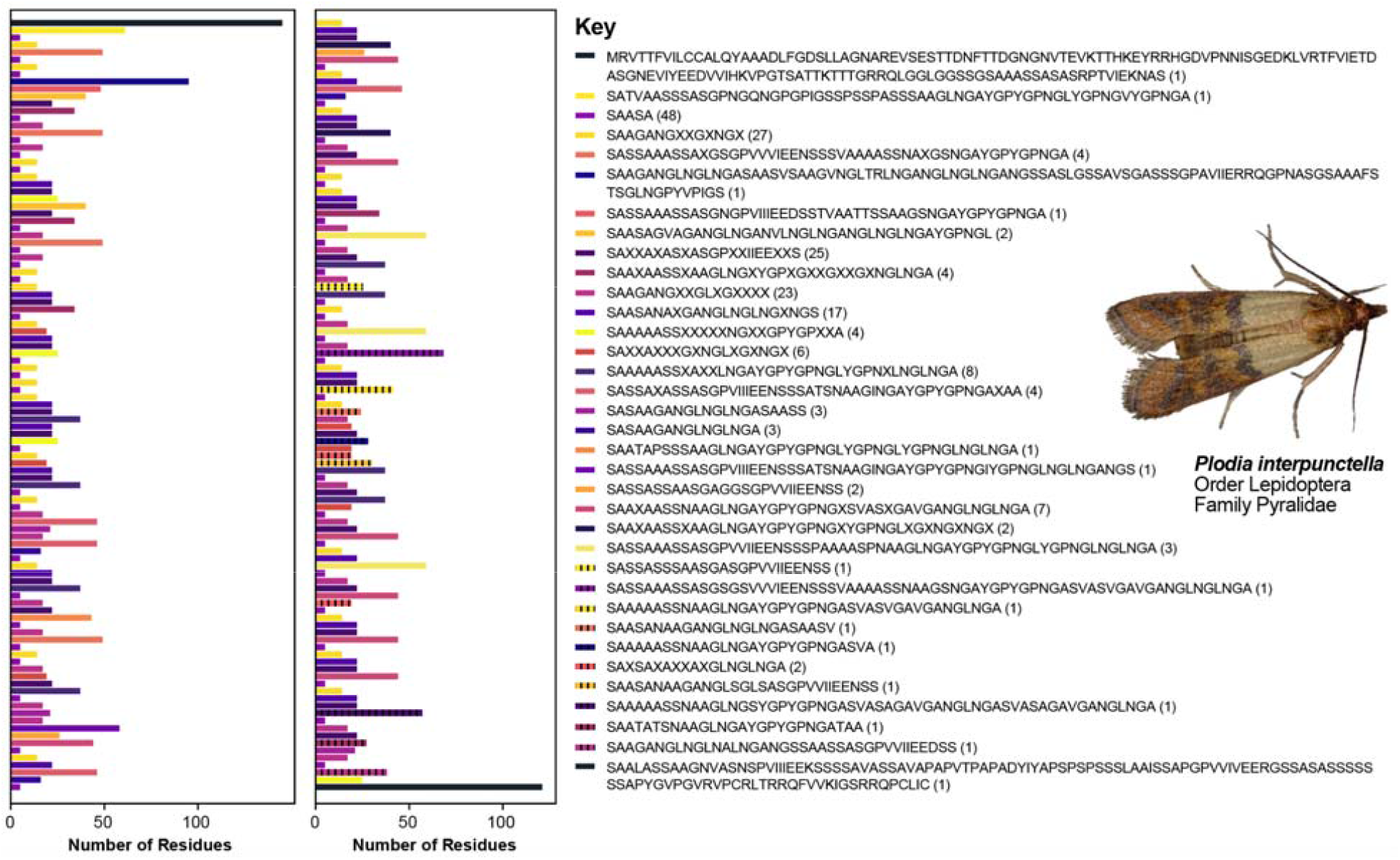
Schematic of the identity and ordering of repeat motifs in *P. interpunctella*. On the right panel are the repetitive units with the *N-*terminus at the beginning and the *C*-terminus at the end. The number in parenthesis refers to the number of times that particular motif is repeated across the gene. The color corresponds with the ordering of the repeats shown on the left. The gene is split into two panels, starting in the left panel and continuing in the right panel. “X” implies a site that is variable.

**Figure 3.**
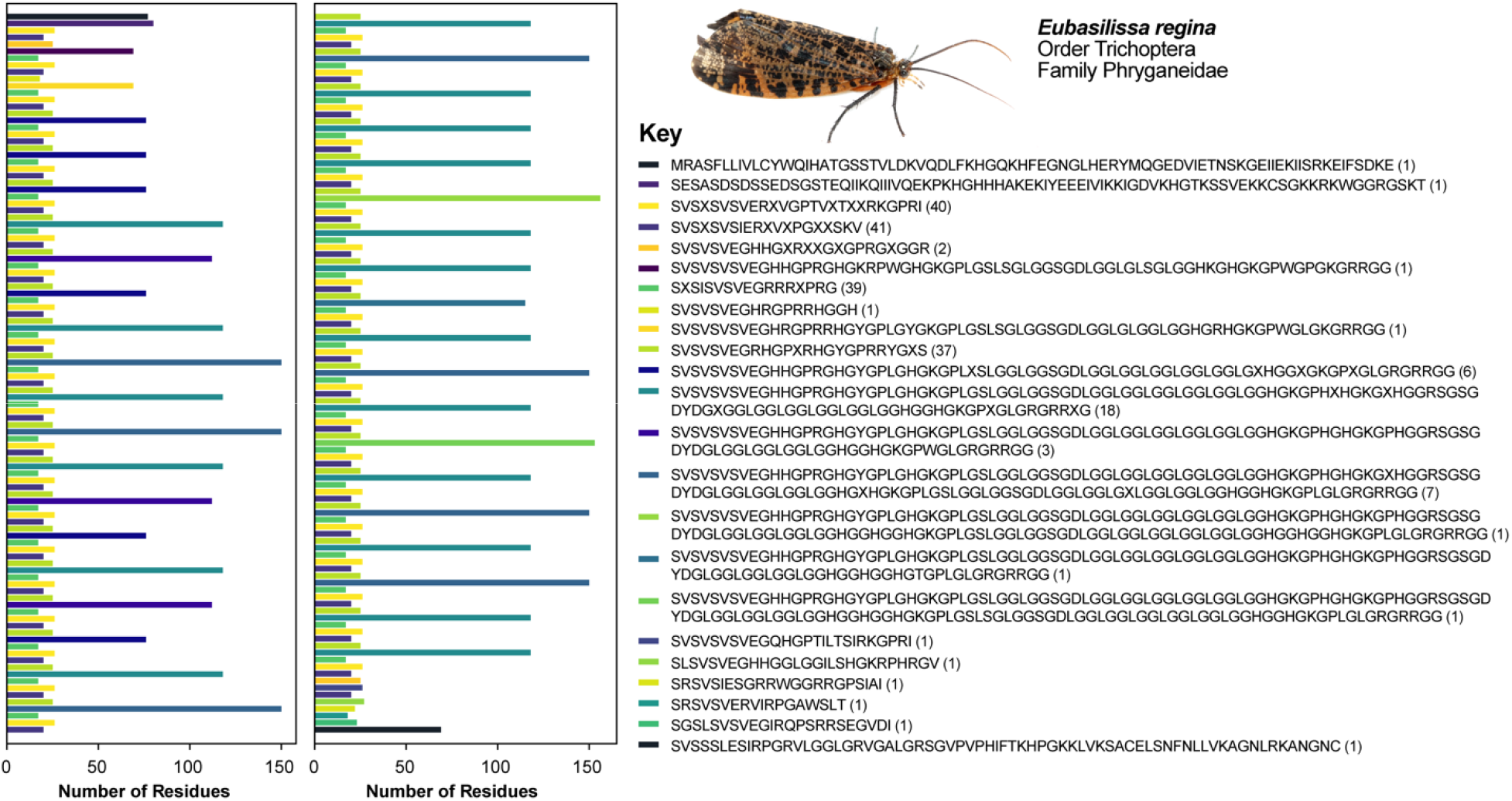
Schematic of the identity and ordering of repeat motifs in *E. regina*. On the right panel are the repetitive units with the *n-*terminus at the beginning and the *c*-terminus at the end. The number in parenthesis refers to the number of times that particular motif is repeated across the gene. The color corresponds with the ordering of the repeats shown on the left. The gene is split into two panels, starting in the left panel and continuing in the right panel.

### Genome annotation

For the structural annotations of the genomes, we masked and annotated repetitive elements using RepeatMasker [29] after identifying and classifying them d*e novo* with RepeatModeler2 [30] following [31]. For species specific gene model training, we used BUSCO v.4.1.4 [21] with the Endopterygota odb10 core ortholog sets [32] with the –long option in genome mode. In addition, we predicted genes with the homology–based gene prediction GeMoMaPipeline of GeMoMa v1.6.4 [33-34] using previously published genomes. For *E. regina* we used the genome of *Agypnia vestita* (JADDOH000000000.1) [35] and for *P. interpunctella* we used the genome of *Bombyx mori* (GCF_014905235) as reference. We then used the MAKER v3.01.03 pipeline [36] to generate additional *ab initio* gene predictions with the proteins predicted from GeMoMa for protein homology evidence and the augustus-generated gene prediction models from BUSCO for gene prediction. For EST evidence, we used the transcriptome of *Ptilostomis semifasciata* (111015_I297_FCD05HRACXX_ L1_INSbttTHRAAPEI-17, 1kite.org) for *E. regina* and Iso-seq data for *P. interpunctella*. Evidence used in Maker and the Maker config files can be found in the *Gigascience* GigaDB repository [26].

To add functional annotations to the predicted proteins, we blasted the predicted proteins against the ncbi-blast protein database using BlastP in blast.2.9 with an e-value cutoff of 10-4 and –max_target_seqs set to 10 (see repository). We then used the command line version of Blast2GO v.1.4.4 [37] to assign functional annotation and GO terms.

### Validation and quality control

In addition to full-length *H-fibroin* sequences, we recovered a high number of single copy orthologs in each genome with BUSCO. The *E. regina* genome contained 95.2% of an Endopterygota core gene collection (comprised of 2124 genes) indicating an almost complete coverage of known single copy orthologs in the coding fraction. While the number of single-copy orthologs recovered in the new *P. interpunctella* genome was similar to earlier published genomes (99.3% of the Endopterygota core gene collection, 99.1% of the Lepidoptera core gene collection), the full-length sequence of *H-fibroin* only recovered in the HiFi based genome gives some indication of how other portions of the genome may have assembled. Following contamination screening by NCBI, we filtered out three instances of *Wolbachia* contamination in the *P. interpunctella* genome. BlobPlots for both genomes revealed low levels of contamination (Supplementary Figures 1, 2).

### Structural and functional annotation

A total of 56.26% of the *E. regina* genome was classified as repetitive (54.2% interspersed repeats). More than half of the interspersed repeats, 29.87%, could not be classified by comparison with known repeat databases, and therefore may be specific for Trichoptera. Of the repeats that were classified, retroelements were the most abundant, comprising 15.35% (of which 14.55% are LINEs) of the genome. The relatively high proportion of repetitive sequence supports previous studies which suggest that repetitive element expansion occurred in lineages of tube case-making caddisflies, such as the closely related genera *Agrypnia* and *Hesperophylax* [9, 35]. In contrast, a total of 31.94% of the *P. interpunctella* genome assembly was masked as repeats. A total of 23.04% of the annotated repeats were interspersed repeats. Details on the repeat classes are given in the *Gigascience* GigaDB repository [26].

The genome annotations resulted in the prediction of 16,937 and 60,686 proteins in *P. interpunctella* and *E. regina*, respectively. Of the annotated proteins, for *E. regina*, 28,358 showed significant sequence similarity to entries in the NCBI nr database, of those 12,550 were mapped to GO terms, and 5,652 were functionally annotated with Blast2GO. For *P. interpunctella*, 16,349 were verified by BLAST, 12,410 were mapped to GO terms, and 9,711 were functionally annotated in Blast2GO.

The major biological process found in the two genomes were cellular (*E. regina*: 2,326 genes; *P. interpunctella:* 4,725 genes) and metabolic (*E. regina*: 2,454 genes; *P. interpunctella:* 3,699 genes) processes. Binding (*E. regina*: 2,382 genes; *P. interpunctella:* 4,405 genes) and catalytic activity (*E. regina*: 2,778 genes; *P. interpunctella:* 3,893 genes) were the largest subcategories in molecular function. Regarding the cellular component category, most genes were assigned to the cell (1,553 genes) and membrane (1,491 genes) subcategory in *E. regina* and to the cellular anatomical entity subcategory in *P. interpunctella* (5,602 genes). The major biological process found in both genomes were cellular and metabolic processes.

### Re-use potential

We provide a complete genome of two species of silk-producing insects in the superorder Amphiesmenoptera, the moth *P. interpunctella* and the caddisfly *E. regina*, and recover the difficult-to-sequence repetitive regions of both genomes with HiFi sequencing. *P. interpunctella* is currently being developed in multiple labs as a model organism and this genome assembly will facilitate molecular genetics research on this species. We show that PacBio HiFi sequencing allows for accurate generation of repetitive protein-coding regions of the genome (silk *fibroins*), and this likely applies to other similarly repetitive regions of the genome. For Trichoptera, there are only four other HiFi genome assemblies available on Genbank, only one of which has been published [38] and insects have generally been neglected (relative to their total species diversity) with respect to genome sequencing efforts [15-16], which is especially true for aquatic insects [14]. These data serve as the first step to study the evolution of adhesive silk in Amphiesmenoptera, which is an innovation that is beneficial for survival in aquatic and terrestrial environments. Finally, the Iso-seq data that we provide serve as useful resources for the translational aspects of silk – these data provide information on how Amphiesmenoptera genetically modulate and regulate different silk properties, that allow them using silk for different purposes such as for nets, cases, and cocoons in both terrestrial and aquatic environments.

## Availability of source code and requirements

All custom-made scripts used in this study are available on GitHub (https://github.com/AshlynPowell/silk-gene-visualization/tree/main).

## Availability of supporting data

Raw sequence data, genome assemblies, and sample information are all available from NCBI (accession can be found in Table 1). All supporting data and materials are available on GigaDB.

## Declarations

All authors have nothing to declare.

## Competing interests

The authors declare that they have no competing interests.

## Funding

Smithsonian National Museum of Natural History Global Genome Initiative (GGI-Peer-2018-182) to T.P.C., R.D., T.D., A.Y.K., Smithsonian Museum Conservation Institute Federal and Trust funds to T.P.C.. and P.B.F. University of Florida Research Opportunity Seed Fund internal award number AWD06265 to PIs AYK and CGS.

## Authors’ contributions

AYK: Designed project, collected samples, provided computational resources, manuscript writing.

CGS: Designed project, data analysis, manuscript writing.

AM: Sample preparation, manage colonies, manuscript writing.

JH: Data analysis, manuscript writing.

AP: Data analysis, manuscript writing.

DP: Data file management, manuscript writing.

SH: Visualization, manuscript writing.

TPC: Grant writing, manuscript writing.

RBD: Grant writing, manuscript writing.

TD: Grant writing, manuscript writing.

RBK: Collected samples, manuscript writing.

RM: Helped with sample preparation, manage colonies, manuscript writing.

SUP: provided computational resources, manuscript writing.

RJS: Grant writing, manuscript writing.

KT: Collected samples, manuscript writing.

PBF: Designed project, collected samples, conducted analyses, provided computational resources, manuscript writing.

## Acknowledgements

We thank the BYU, UF and LOEWE-Centre for Translational Biodiversity Genomics (TBG) High Performance Computing clusters for providing the computational resources needed to complete this study. TBG is funded by the Hessen State Ministry of Higher Education, Research and the Arts (HMWK), which financially supported J.H. and S.U.P.. S.H. was supported by NSF award #OPP-1906015.

## Authors’ information

AM: Graduate Student at the University of Florida, School of Natural Resources and Environment. Studies variation in lepidopteran silk production.

AP: Undergraduate Student at Brigham Young University. Studies the genomics of caddisfly silk.

AYK: Associate Curator at the University of Florida. Works on Lepidoptera, genomics, and evolution.

CGS: formally Assistant Scientist at the University of Florida. Expert on silks and Lepidoptera. Currently works at Pacific Biosciences.

DP: Project Manager at the Florida Museum of Natural History, University of Florida. Works on Lepidoptera systematics and evolution.

JH: Postdoctoral researcher at LOEWE TBG, interested in biodiversity genomics, comparative genomics, evolution and phylogenomics.

KT: Researcher at Shinshu University, Japan, specialist of Trichoptera.

PBF: Assistant Professor of Genetics, Genomics, and Biotechnology at Brigham Young University, specialist on the genomics of caddisflies and their silk.

RBD: Data Scientist at Smithsonian National Museum, Washington, D.C.

RJS: Professor of Bioengineering, University of Utah, specialist on the biomechanics of caddisfly silk.

RBK: Guest Professor of Kanagawa Institute of Technology, specialist of Trichoptera.

RM: Biological Scientist at the University of Florida researching molecular biology and genetics of organisms.

SH: Postdoctoral Research Associate at Washington State University and a specialist in insect genomics.

SUP: Entomologist at Senckenberg Research Institute and Natural History Museum Frankfurt and Justus-Liebig-University, interested in the evolution and ecology of freshwater insects, especially Trichoptera.

TD: Researcher and curator at Smithsonian National Museum of Natural History, Washington, D.C., Works on biodiversity of flies and is curator for aquatic insects.

TPC: Physical Scientist at the Smithsonian Museum Conservation Institute.

## Supplementary Files

**Supplementary Figure 1(in GigaDB).**
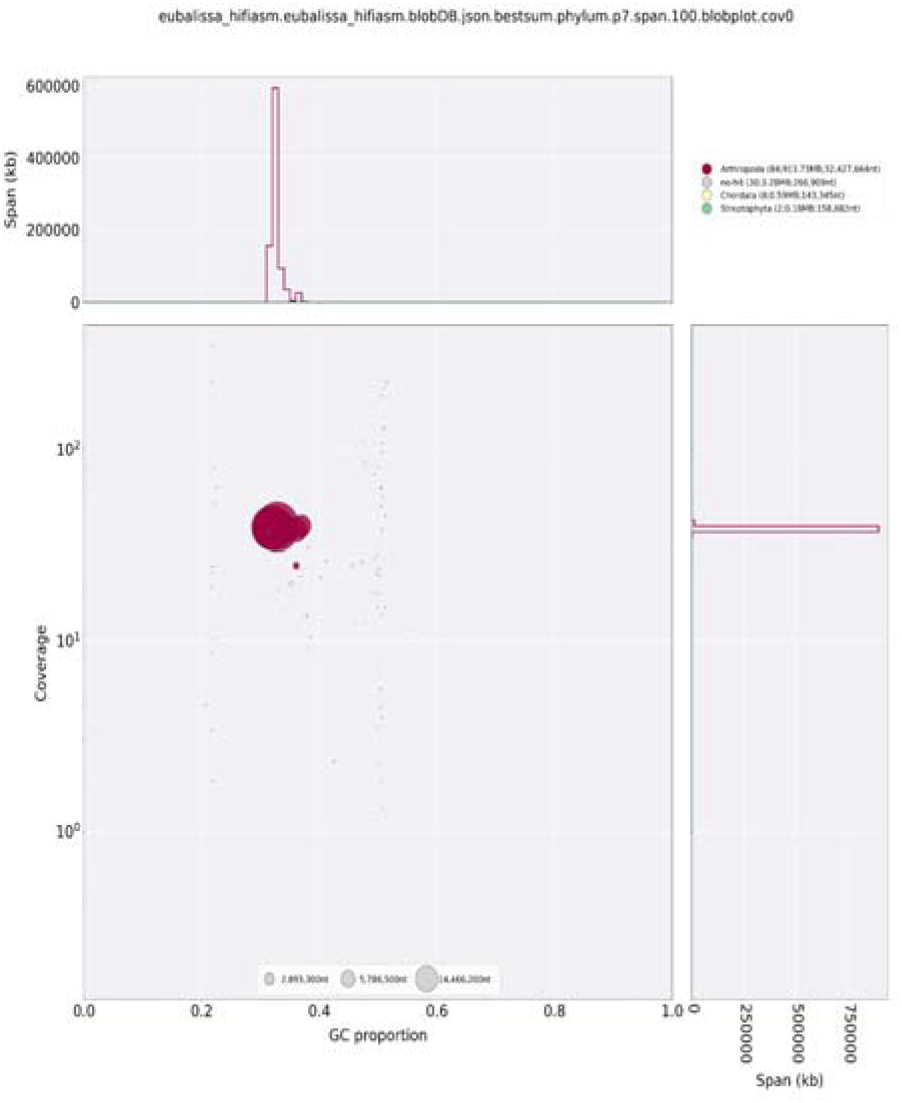
BlobPlot for *E. regina*.

**Supplementary Figure 2(in GigaDB).**
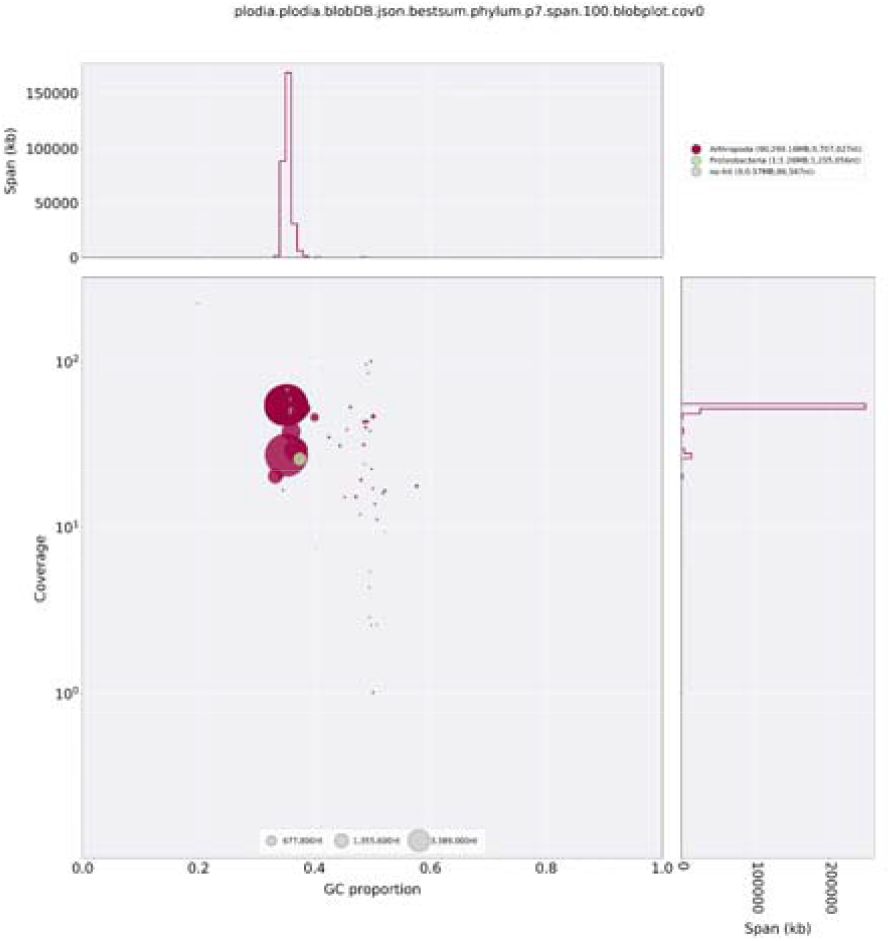
BlobPlot for *P. interpunctella*.

## Notes

### Competing Interest Statement

The authors have declared no competing interest.

## References

1 Numata K. How to define and study structural proteins as biopolymer materials. Polym. J., 2020; 52: 1043–1056. doi:10.1038/s41428-020-0362-5.

2 Davies PL, Hew CL. Biochemistry of fish antifreeze proteins. FASEB J, 1990; 4: 2460–2468. doi:10.1096/fasebj.4.8.2185972

3 Kono N, Nakamura H, Ohtoshi R et al. The bagworm genome reveals a unique fibroin gene that provides high tensile strength. Commun. Biol., 2019; 2: 1–9. doi:10.1038/s42003-019-0412-8.

4 Stewart RJ, Wang CS. Adaptation of caddisfly larval silks to aquatic habitats by phosphorylation of H-fibroin serines. Biomacromolecules, 2010; 11: 969–974. doi:10.1021/bm901426d.

5 Ashton NN, Roe DR, Weiss RB et al. Self-tensioning aquatic caddisfly silk: Ca2+-dependent structure, strength, and load cycle hysteresis. Biomacromolecules, 2013; 14: 3668–3681. doi:10.1021/bm401036z.

6 You Z, Ye X, Ye L et al. Extraordinary mechanical properties of composite silk through hereditable transgenic silkworm expressing recombinant major ampullate spidroin. Sci. Rep., 2018; 8: 1–4. doi:10.1038/s41598-018-34150-y.

7 Sutherland TD, Young JH, Weisman S et al. Insect silk: one name, many materials. Annu. Rev. Entomol., 2010; 55: 171–188. doi:10.1146/annurev-ento-112408-085401.

8 Zhou CZ, Confalonieri F, Medina N et al. Fine organization of Bombyx mori fibroin heavy chain gene. Nucleic Acids Res., 2000; 28: 2413–2419. doi:10.1093/nar/28.12.2413.

9 Heckenhauer J, Frandsen PB, Sproul JS et al. Genome size evolution in the diverse insect order Trichoptera. GigaScience, 2022; 11: giac011. doi:10.1093/gigascience/giac011.

10 Yonemura N, Mita K, Tamura T et al. Conservation of silk genes in Trichoptera and Lepidoptera. J. Mol. Evol., 2009; 68: 641–653. doi:10.1007/s00239-009-9234-5.

11 Rutschky CW, Calvin D. Indian meal moth. 1990; https://extension.psu.edu/indian-meal-moth. Accessed March 2022.

12 Fasulo TR, Knox MA. Indianmeal moth - Plodia interpunctella (Hübner). 1998; https://entnemdept.ufl.edu/creatures/urban/stored/indianmeal_moth.HTM. Accessed March 2022.

13 Wiggins, G. B. The caddisfly family Phryganeidae (Trichoptera)., 1998; University of Toronto Press, Toronto, Buffalo, London.

14 Hotaling S, Kelley JL, Frandsen PB. Aquatic insects are dramatically underrepresented in genomic research. Insects, 2020; 11: 601. doi:10.3390/insects11090601.

15 Hotaling S, Sproul JS, Heckenhauer J et al. Long reads are revolutionizing 20 years of insect genome sequencing. Genome Biol. Evol., 2021; 13: evab138. doi:10.1093/gbe/evab138.

16 Hotaling S, Kelley JL, Frandsen PB. Toward a genome sequence for every animal: Where are we now? P. Natl. Acad. Sci. USA, 2021; 118: e2109019118. doi:10.1073/pnas.2109019118.

17 Kokot M, Długosz M, Deorowicz S. KMC 3: counting and manipulating k-mer statistics. Bioinformatics, 2017; 33: 2759–2761. doi:10.1093/bioinformatics/btx304.

18 Ranallo-Benavidez TR, Jaron KS, Schatz MC. GenomeScope 2.0 and Smudgeplot for reference-free profiling of polyploid genomes. Nat. Commun., 2020; 11: 1432. doi:10.1038/s41467-020-14998-3.

19 Cheng H, Concepcion GT, Feng X et al. Haplotype-resolved de novo assembly using phased assembly graphs with hifiasm. Nat. Methods, 2021; 18: 170–175. doi:10.1038/s41592-020-01056-5.

20 Trizna M. assembly_stats 0.1.4 | Zenodo. 2020; https://zenodo.org/record/3968775#.Yi_b4C1h2FU. Accessed March 2022. doi:10.5281/zenodo.3968775.svg.

21 Manni M, Berkeley MR, Seppey M et al. BUSCO update: novel and streamlined workflows along with broader and deeper phylogenetic coverage for scoring of eukaryotic, prokaryotic, and viral genomes. Mol. Biol. Evol., 2021; 38: 4647–4654. doi:10.1093/molbev/msab199.

22 Laetsch DR, Blaxter ML. BlobTools: Interrogation of genome assemblies [version 1; peer review: 2 approved with reservations]. F1000Research, 2017; 6: 1287. doi:10.12688/f1000research.12232.1.

23 Li H. Minimap2: pairwise alignment for nucleotide sequences. Bioinformatics, 2018; 34: 3094–3100. doi:10.1093/bioinformatics/bty191.

24 Kludkiewicz B, Kucerova L, Konikova T et al. The expansion of genes encoding soluble silk components in the greater wax moth, Galleria mellonella. Insect Biochem. Molec., 2019; 106: 28–38. doi:10.1016/j.ibmb.2018.11.003.

25 Stanke M, Diekhans M, Baertsch R et al. Using native and syntenically mapped cDNA alignments to improve de novo gene finding. Bioinformatics, 2008; 24: 637–644. doi:10.1093/bioinformatics/btn013.

26 Kawahara AY, Storer CG, Markee A et al. Supporting data for “Long-read HiFi sequencing correctly assembles repetitive heavy fibroin silk genes in new moth and caddisfly 17 genomes”. GigaScience Database. 2022. doi:XXXX. [Repository for supplemental files for this paper]

27 Luo S, Tang M, Frandsen PB et al. The genome of an underwater architect, the caddisfly Stenopsyche tienmushanensisHwang (Insecta: Trichoptera). GigaScience, 2018; 7: giy143. doi:10.1093/gigascience/giy143.

28 Frandsen PB, Bursell MG, Taylor AM et al. Exploring the underwater silken architectures of caddisworms: comparative silkomics across two caddisfly suborders. Philos. T. R. Soc. B, 2019; 374: 20190206. doi:10.1098/rstb.2019.0206.

29 Smit AFA, Hubley R, Green P. RepeatMasker Open-4.0. 2013–2015; http://www.repeatmasker.org. Accessed January 2022.

30 Flynn JM, Hubley R, Goubert C et al. RepeatModeler2 for automated genomic discovery of transposable element families. P. Natl. Acad. Sci. USA, 2020; 117: 9451–9457. doi:10.1073/pnas.1921046117.

31 Heckenhauer J, Frandsen PB, Gupta DK et al. Annotated draft genomes of two caddisfly species Plectrocnemia conspersa CURTIS and Hydropsyche tenuis NAVAS (Insecta: Trichoptera). Genome Biol. Evol., 2019; 11: 3445–3451. doi:10.1093/gbe/evz264.

32 Kriventseva EV, Kuznetsov D, Tegenfeldt F et al. OrthoDB v10: sampling the diversity of animal, plant, fungal, protist, bacterial and viral genomes for evolutionary and functional annotations of orthologs. Nucleic Acids Res., 2019; 47: D807–811. doi:10.1093/nar/gky1053

33 Keilwagen J, Wenk M, Erickson JL et al. Using intron position conservation for homology-based gene prediction. Nucleic Acids Res., 2016; 44: e89. doi:10.1093/nar/gkw092.

34 Keilwagen J, Hartung F, Paulini M et al. Combining RNA-seq data and homology-based gene prediction for plants, animals and fungi. BMC Bioinformat., 2018; 19: 1–2. doi:10.1186/s12859-018-2203-5.

35 Olsen LK, Heckenhauer J, Sproul JS et al. Draft genome assemblies and annotations of Agrypnia vestita Walker, and Hesperophylax magnus Banks reveal substantial repetitive element expansion in tube case-making caddisflies (Insecta: Trichoptera). Genome Biol. Evol., 2021; 13: evab013. doi:10.1093/gbe/evab013.

36 Campbell MS, Holt C, Moore B et al. Genome annotation and curation using MAKER and MAKERLP. Curr. Protocols Bioinformat. 2014; 48: 4–11. doi:10.1002/0471250953.bi0411s48.

37 Conesa A, Götz S. Blast2GO: a comprehensive suite for functional analysis in plant genomics. Int. J. Plant Genomics. 2008; 2008: 1–12. doi:10.1155/2008/619832.

38 Ríos-Touma B, Holzenthal RW, Rázuri-Gonzales E et al. De novo genome assembly and annotation of an Andean caddisfly, Atopsyche davidsoni Sykora, 1991, a model for genome research of high-elevation adaptations. Genome Biol. Evol., 2022; 14: evab286. doi:10.1093/gbe/evab286.

